# Recombinant human MG53 protein preserves mitochondria integrity in cardiomyocytes during ischemia reperfusion-induced oxidative stress

**DOI:** 10.1101/2020.02.06.936278

**Authors:** Kristyn Gumpper, Hanley Ma, Karthikeyan Krishnamurthy, Xinyu Zhou, Ki Ho Park, Matthew Sermersheim, Jingsong Zhou, Tao Tan, Pei-Hui Lin, Lei Li, Jianxun Liu, Hua Zhu, Jianjie Ma

**Affiliations:** The Ohio State University Wexner Medical Center Department of Surgery, Columbus, OH 43210; Department of Biochemistry, Brown University, Providence, RI 02912; University of Texas at Arlington Department of Kinesiology, Arlington, TX 76109; Xiyuan Hospital, China Academy of Chinese Medical Sciences, Beijing 100091, China

**Author notes:** Corresponding Author: Jianjie Ma, PhD, 460 W 12^th^ Avenue, Columbus, OH 43210.

**Keywords:** cardioprotection, mitophagy, cell membrane repair, TRIM72, heart failure

## Abstract

Ischemic injury to the heart causes a loss of mitochondria function due to an increase in oxidative stress. MG53, also known as TRIM72, is highly expressed in striated muscle and is essential to repair damage to plasma membrane. We have shown that *mg53-/-* mice are more susceptible to ischemia-reperfusion injury, whereas treatment with exogenous recombinant human MG53 (rhMG53) reduces both infarct damage and restores cardiac function. This study assesses whether MG53 protects and repairs mitochondria injury after oxidative stress associated with myocardial infarction. We hypothesize that in addition to known cell membrane repair function, MG53 acts as a myokine to protect cardiomyocytes by maintaining mitochondrial function. A combination of *in vivo* and *in vitro* ischemia/reperfusion models were used to assess MG53’s effect on mitochondria using biochemical assays and confocal microscopic imaging. Treatment with rhMG53 allowed cells to maintain a healthy mitochondrial membrane potential, reduced release of mitochondrial reactive oxygen species, and mitigated mitophagy. Mitochondrial localization of rhMG53 is mediated by exposure of and interaction with cardiolipin on the mitochondrial membrane. Our data demonstrates that rhMG53 protein preserves mitochondria integrity in cardiomyocytes during ischemia reperfusion-induced oxidative stress.

## Introduction

Ischemic heart disease remains one of the top causes of mortality world-wide [1]. Ischemic injury causes a loss of healthy mitochondria, resulting in reduced energy production and death of cardiomyocytes [2]. During an ischemia and reperfusion event like a myocardial infarction (MI), cardiomyocytes experience mitochondria dysfunction and the release of reactive oxygen species (ROS) from the mitochondria that further damages the cell [3]. Therapeutic approaches that can preserve mitochondria integrity will be valuable to improving survival after MI.

MG53, also known as TRIM72, is highly expressed in skeletal muscle and modestly in other tissues. It is essential for repairing damage to the plasma membrane [4] and can be secreted from skeletal muscle where it acts as a myokine to repair other damaged tissues [5]. When cells are damaged, there is an increase in oxidative stress and the plasma membrane inner leaflet lipid phosphatidylserine (PS) is exposed to signal for membrane repair. MG53’s membrane repair function is associated with changes in oxidative state inside the cell [4, 6], and extracellular MG53 can target the exposed PS to facilitate repair of injured tissues [7]. We have shown that *mg53*−*/*− mice are more susceptible to injury, and treatment with exogenous recombinant human MG53 (rhMG53) reduces damage and restores functions in tissues like heart [8, 9], kidney [10], and lung [11].

To date, MG53’s link to mitochondrial protection is tenuous at best. Chung et al [12] found enrichment of TRIM72 (MG53) in the mitochondria fraction as part of the cardioprotective proteins in mouse heart subjected to ischemia/reperfusion injury. In a study on metabolic syndrome, Ma et al [13] demonstrated MG53 co-localization with mitochondria in skeletal muscles derived from mice fed with high fat diet. Lijie et al [14] showed that the expression of MG53 and Ambra1 (activating molecule in Beclin-1 regulated autophagy) in skeletal muscle derived from rats with chronic kidney disease was decreased, suggesting a potential link between MG53 and mitophagy, a biological process that provides quality control for mitochondria. While these studies indicate a potential interaction of MG53 with the mitochondria, the mechanism behind MG53’s role in facilitating mitochondrial protection remains unknown.

In this study, we test the hypothesis that MG53 can either directly prevent oxidative stress-induced injury to mitochondria, or indirectly preserve the pool of healthy mitochondria through modulation of mitophagy. We found that MG53 can bind to cardiolipin (CL), a mitochondria specific phospholipid, to prevent injury to the mitochondria. We further showed that rhMG53 treatment mitigates the engulfment of mitochondria by the lysosome. Thus, MG53’s role in preserving the mitochondria integrity may reduce the necessity of mitophagy during the acute phase of MI.

## Methods

### Porcine model of angioplasty-induced myocardial infarction

Chinese experimental miniature swine were provided by Beijing Experimental Animal Reproduction and Regulation Center (Grade II, Certificate No. Jing-030). All porcine experiments in this study were performed in accordance with China Academy of Chinese Medical Sciences Guide for Laboratory Animals that conforms to the Guide for the Care and Use of Laboratory Animals published by the U.S. National Institutes of Health. Experimental pigs underwent balloon inflation of the left anterior descending (LAD) coronary artery according to established methods as described [8]. rhMG53 was administered at different times of experimental intervention through the jugular vein.

### Murine model of myocardial infarction

All murine experiments in this study were performed in accordance with The Ohio State University IACUC-approved protocols. The *mg53-/-* and wild type littermate mice were used. Saline or 1 mg/kg rhMG53 dissolved in saline was injected via the tail vein 10 minutes prior to induction of myocardial infarction via left anterior descending (LAD) coronary artery ligation. Ischemia was maintained for 60 minutes. After ischemia, the mouse recovered for 6 hours before cardiac tissue collection for histology and immunofluorescent staining.

### Confocal microscopy

Porcine and mouse tissue samples were embedded in paraffin, sectioned (5 μm thickness), and stained with antibodies against MG53 [4] and COX IV (Cell Signaling Technologies, 11967S). MG53 (633 nm), COX IV (546 nm) and DAPI (405nm) stained slides were imaged on a Zeiss 780 Confocal microscope using Zen 2012 software (Zeiss). Co-localization and fluorescent intensity was quantified using FIJI (ImageJ). HL-1 cells cultured on 35 mm glass bottom dishes were incubated with either 50 nM TMRE or 5 μM MitoSOX Red (ThermoFisher, M36008) for 15 min at 37°C protected from light. Cells were rinsed and imaged in BSS (140 mM NaCl, 2.8 mM KCl, 2 mM MgCl_2_, 1 mM CaCl_2_, 12 mM Glucose, 10 mM HEPES, pH 7.4). Images were collected with a Zeiss 780 confocal microscope and analyzed in FIJI, measuring the fluorescent integrated density of each cell.

### Mitochondrial FLASH event measurement in isolated mouse cardiomyocytes

Transgenic mice with expression of mt-cpYFP provides assessment of the dynamic changes in superoxide signal from the mitochondria [15-20] Cardiomyocytes were isolated from the mt-cpYFP transgenic mice following the protocol of Wang et al [17]. The isolated cardiomyocytes were subjected to 3 hours of hypoxia and 2 hours of reoxygenation. Live cell imaging of the dynamic changes of the mt-cpYFP fluorescence was conducted on a Zeiss 510 confocal microscope at 40x at 1 frame/second by alternating excitation at 488 nm and collecting emission at 561 nm [17]. Flash events were analyzed using FIJI (ImageJ) [21].

### Oxidative damage to HL-1 cardiomyocytes

HL-1 cardiomyocytes (Sigma, SCC065) were cultured at low passages (7-20) according to established protocols [22, 23]. For H_2_O_2_ damage, HL-1 cells were subjected to 300 μM H_2_O_2_ in unsupplemented Claycomb media (Sigma, 51800C) for 1 hour and recovered with either 10 µg/mL BSA or rhMG53 for 2 hours. To generate a hypoxia/reoxygenation model of oxidative stress, we adapted the protocol for hypoxia, energy depletion, and acidosis (HEDA) from Åström-Olsson et al [24]. Briefly, HL-1 cells in PBS pH 6.7 without glucose, Mg^2+^ or Ca^2+^ were placed in a sealed chamber gassed with 5% CO_2_, 1% O_2_, and 94% N_2_ to establish the HEDA environment for 24 hours. After 24 hours, cells were reoxygenated in BSS at 37°C for 1 h. Control cells were incubated in BSS in a tissue culture incubation chamber at 5% CO_2_ and 37°C. For both conditions, cells were incubated with either 10 µg/mL BSA or rhMG53.

### Lipid binding assays

ELISA Snoopers® (Avanti Polar Lipids) lipid strips were used to assess rhMG53 binding capacity to PS and CL. 8-well strips pre-coated with either lipid were incubated with increasing doses of rhMG53 (0, 0.3125, 0.625, 1.25, 2.5, 5, and 10 µg/mL) ELISAs were performed according to manufacturer’s protocol. Monoclonal MG53 antibody (5259 generated by Dr. Hiroshi Takeshima, Kyoto, Japan) [4] conjugated to biotin was used for detection. Signal was developed using Streptavidin-HRP detection antibody and TMB Substrate Reagent (BD OptEIA, 555214) and quantified using a Flex Station III plate reader.

### Mitophagy assay

pCHAC-mt-mKeima plasmid (Addgene plasmid #72342) [25], was packaged into lentivirus and infected into HL-1 cells. Cells were imaged on a Nikon microscope in a 37°C heated and humidified chamber gassed with 5% CO_2_ in HL-1 media. Movies were taken on a Nikon A1R excitation at 488 and 561 reading at an emission of 560-720 for 2 hours once 1mM H_2_O_2_ was added to the media. Images were collected immediately and 2 hours later. Images were analyzed for fluorescent intensity for the 488 excitation channel (green) and the 561 excitation channel (red) using FIJI (ImageJ) [21].

### Western blotting

Mitochondria fraction of HL-1 cells was isolated using the Mitochondria Isolation Kit for Cultured Cells (ThermoFisher, 89874) following manufacturer’s protocol. Briefly, HL-1 cells that underwent BSS or HEDA treatment with 10 µg/mL BSA or rhMG53 were trypsinized with TrypLE (ThermoFisher, 12604021) and resuspended in 2 mL Claycomb media. Cells were counted and 8 million cells were collected for fractionation. The remaining cells were lysed with RIPA buffer (10 mM Tris-HCl, pH 7.2, 150 mM NaCl, 1% NP-40, 0.5% SDS, and 0.5% deoxycolate), and supplemented with a cocktail of protease inhibitors (Sigma, S8820) for a total comparison. The 8 million cells were centrifuged to collect the pellet and gently lysed using kit reagents. The supernatant cytosolic fraction was retained from the fractionation. The resulting mitochondrial pellet was lysed with 2% CHAPS in tris-buffered saline.

Lysates were separated on a 10% SDS-PAGE acrylamide gel and transferred to polyvinylidene fluoride membranes (PVDF) (Millipore). The blots were washed with Tris-buffered saline Tween-20 (TBS-T), blocked with 5% milk in TBS-T for 1 hour. Blots were probed with antibodies against MG53, GAPDH (CST, 2118S), and COX IV (CST, 11967S). COX IV was used as a marker for mitochondria enrichment, while GAPDH marked the cytosol fraction. The PVDF membranes were developed by chemiluminescence using SuperSignal west femto maximum sensitivity substrate (Thermo Scientific, 34096) on a ChemiDoc MP Imaging System (Bio-Rad). Bio-Rad Image Lab™Version 6.0 software was used to calculate the band intensity for each western blot.

### Statistical analysis

Data was analyzed by several statistical methods (e.g., paired or unpaired *t*-tests, ANOVA, etc.) using commercial Prism software (GraphPad Prism 8). Data are presented as means ± SD. For comparisons between two groups, significance was determined by Student *t*-test parametric analysis. For comparison of multiple groups, multifactorial analysis of variance (ANOVA) was used to determine statistical significance. Corrections, such as Brown-Forsythe or Welch’s correction, were used if there was a difference in variance between groups. For all statistical tests: *p<0.05, **p<0.01, ***p<0.001, ****p<0.0001

## Results

### rhMG53 targets mitochondria in mouse and pig models of myocardial infarction

In cardiac tissue, MG53 typically localizes to the intercalated discs, an area of constant low-level plasma membrane stretch and injury as the heart expands and contracts for each beat. As shown in **Figure 1A**, antibody against MG53 identified MG53 at the intercalated discs of hearts derived from healthy wild type mice, but not from healthy *mg53-/-* mice. Following MI in *mg53-/-* mice, rhMG53 was administered (tail vein, 1 mg/kg). IHC staining revealed rhMG53 localization to the plasma membrane and intercalated discs (red). Interestingly, rhMG53 also co-localized with the mitochondria, indicated by COX IV signal (green) (**Figure 1B**).

**Figure 1:**
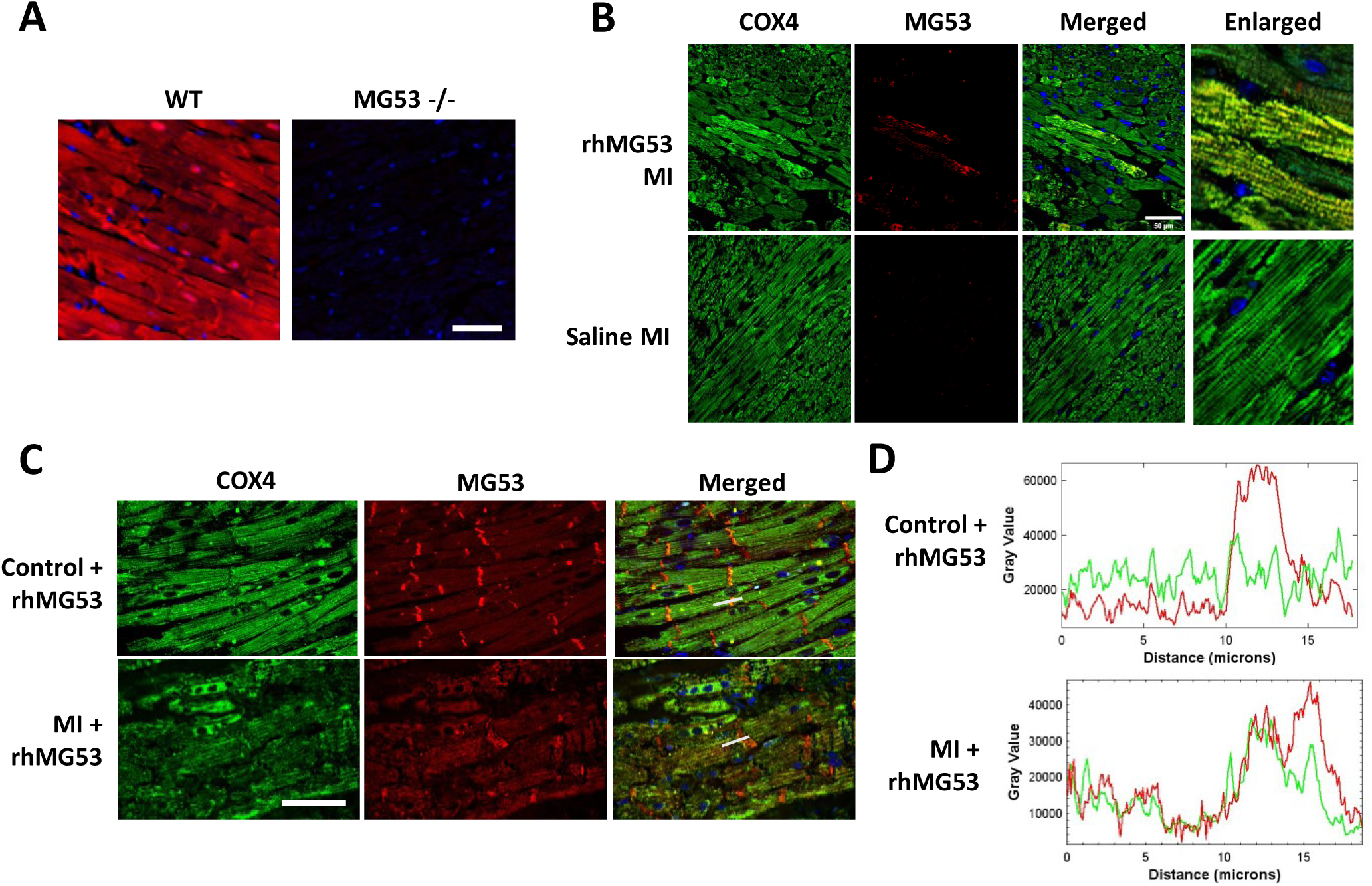
rhMG53 localizes to the intercalated disc and mitochondria in mouse and pig cardiomyocytes after MI. (**A**) IHC reveals MG53 localization at the intercalated disc of WT mice with no staining in the *mg53 -/-* mice. (**B**) IHC of *mg53 -/-* mouse hearts after MI injury revealed no MG53 signal in the saline treatment. Treatment with 2 mg/kg rhMG53 and staining for MG53 (red), COX IV (green), and DAPI (blue) reveals co-localization of MG53 with mitochondria. (**C**) IHC of pig hearts treated with 2 mg/kg rhMG53 with MI injury highlights MG53 localization at the intercalated discs with a translocation to the mitochondria after MI injury. (**D**) Intensity analysis along the indicated grey line in part C shows an over-lapping pattern of MG53 and COX IV in pig heart with MI injury. White scale bar = 50 μm

We confirmed the dual plasma membrane and mitochondria localization in a porcine model of MI (**Figure 1C**). In a healthy pig heart, MG53 localizes mainly to the intercalated disks. However, after a MI, MG53 shifts from the intercalated discs to co-localization with mitochondria. This shift is highlighted in the longitudinal fluorescent intensity analysis in **Figure 1D**, where MG53’s signal is high at the intercalated disc and minimal everywhere else. After a MI, MG53’s intensity at the intercalated disc drops and aligns with the COX IV mitochondrial signal.

This co-localization is consistent with earlier studies in the heart by Chung et al. [12] who found enrichment of TRIM72/MG53 in mitochondria fraction derived from ischemia-reperfusion injured mouse heart. Moreover, the mitochondria localization pattern of rhMG53 in the porcine heart is consistent with our previous observation with skeletal muscle derived from high fat diet treated mice[13].

### rhMG53 reduces superoxide release in isolated cardiomyocytes after hypoxia /reoxygenation

Wang et al. [26] developed a novel indicator targeted to the mitochondrial matrix called circularly permuted yellow fluorescent protein (mt-cpYFP) that selectively fluoresce in the presence of O_2_^−^, the main ROS created by the electron transport chain in the mitochondria. We have recently used this tool to study mitochondrial ROS generation in the mouse model of amyotrophic lateral sclerosis (ALS), termed “mitoFlash” [15, 18-20].

In cardiomyocytes derived from the mt-cpYFP mice, we monitored mitochondrial superoxide generation after hypoxia/reoxygenation following our previous protocol [17] (**Figure 2A**). Under basal conditions, mitoflash events are rare (**Figure 2B**). However, subjecting the cardiomyocytes to hypoxia and reoxygenation significantly increases the frequency of superoxide flashes. When cardiomyocytes are treated with 2 µg/mL rhMG53 before hypoxia/reoxygenation, fewer and smaller superoxide flashes occur. This suggests rhMG53 treatment reduces superoxide production in the mitochondria after oxidative injury.

**Figure 2:**
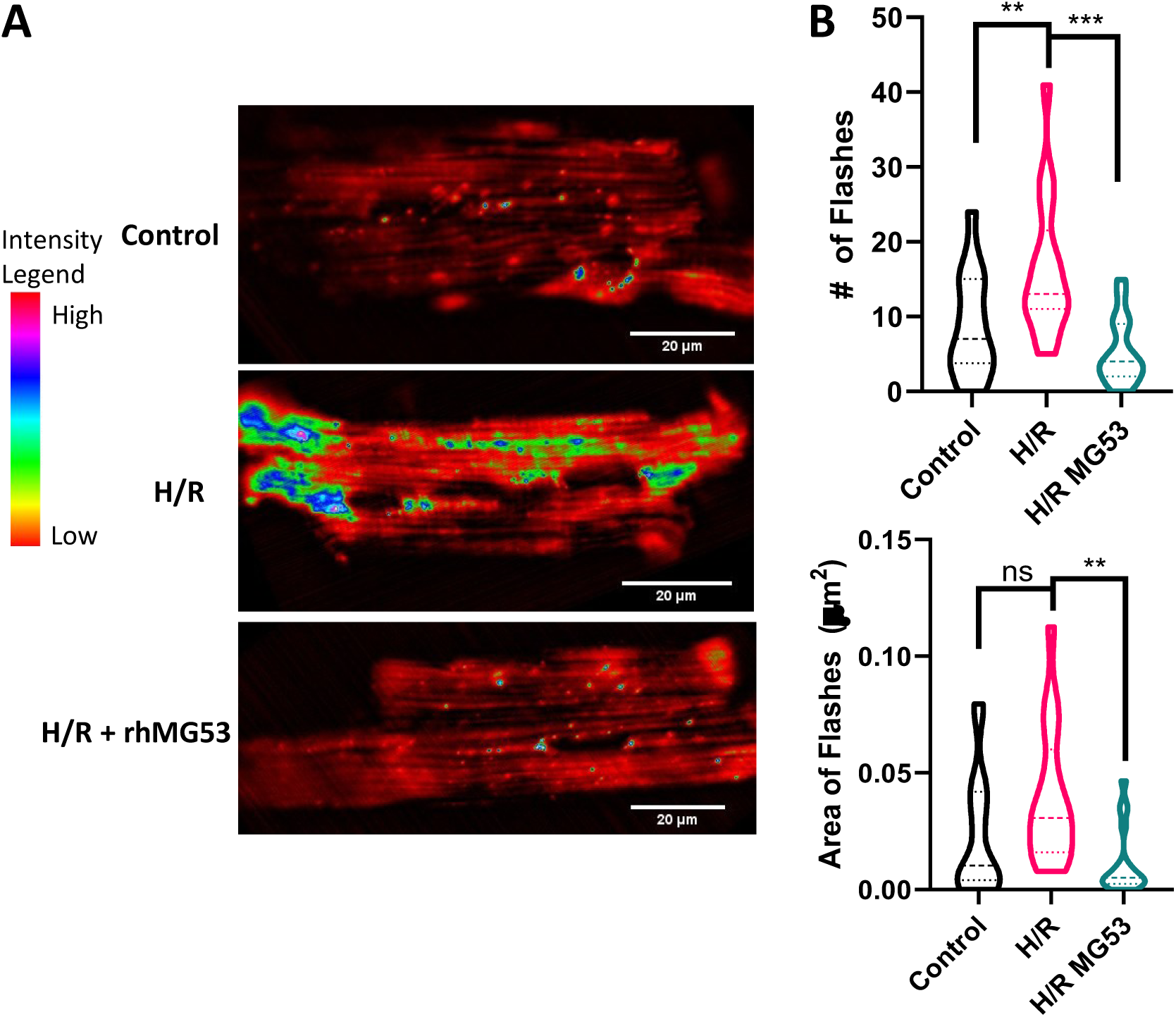
rhMG53 reduces hypoxia/reoxygenation-induced mito-FLASH in adult mouse cardiomyocytes. (**A**) Surface plots of the amplitude and distribution of mitoflashes in cardiomyocytes isolated from mt-cpYFP transgenic mice, under normoxia (control) condition, after hypoxia/reoxygenation (H/R), or hypoxia/reoxygenation +10μg/mL rhMG53 (H/R+rhMG53). (**B**) Hypoxia/reoxygenation increased the number and size of mitoflashes, while treatment with MG53 significantly reduced mitoflash number and size. N=18 Control, 17 H/R, and 17 H/R rhMG53

### rhMG53 preserves mitochondria in HL-1 cells after oxidative stress

Since rhMG53 treatment reduced superoxide release within the mitochondria, we wanted to assess whether rhMG53 also prevented the release of superoxides generated by the mitochondria into the cytoplasm. We used the HL-1 cardiac muscle cell line derived from mice in conjunction with the MitoSOX Red fluorescent indicator to measure rhMG53’s effect on mitochondria after oxidative damage. As shown in **Fig. 3A**, treatment rhMG53 under normoxic conditions did not alter release of superoxides from the mitochondria. However, we observed a significant increase in MitoSOX Red fluorescent intensity in cells that underwent HEDA in the presence of BSA, indicating high concentrations of mitochondrial ROS in the cytoplasm. Treating cells with rhMG53 during HEDA injury resulted in a significantly reduced MitoSOX Red fluorescent intensity like the control conditions (**Figure 3B**). This suggests rhMG53 treatment prevents the release of mitochondria superoxides into the cytoplasm.

**Figure 3:**
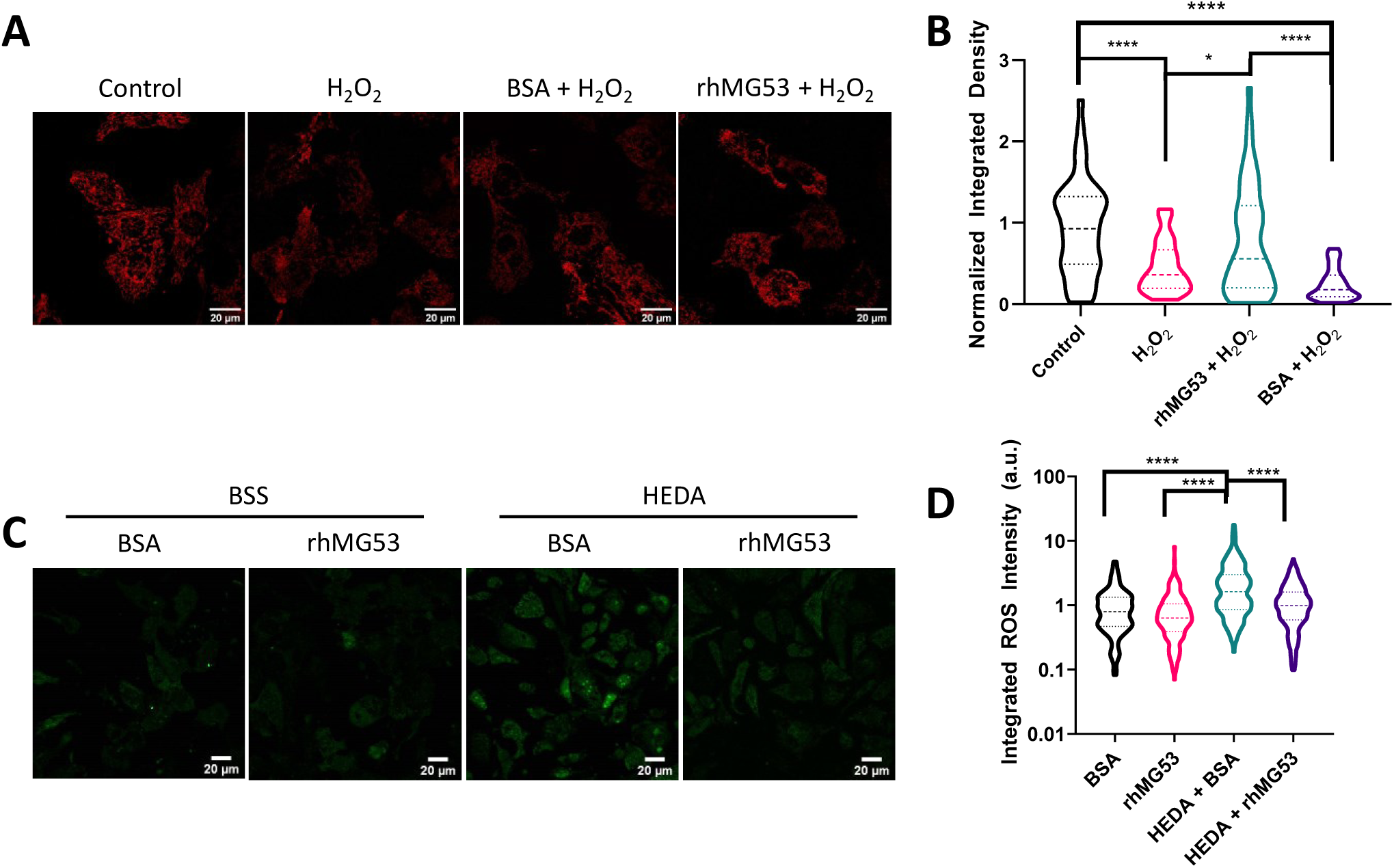
rhMG53 preserves mitochondrial membrane potential and reduces mitochondria-superoxide release in HL-1 cells. (**A**) Treatment of HL-1 cells with 300 µM H2O2 reduces TMRE fluorescence. Compared with BSA, rhMG53 preserved TMRE fluorescence in H2O2-treated HL-1 cells. (**B**) TMRE integrated density for each cell was quantified and normalized to the average intensity of the control cells. N= 46 Control, 37 H_2_O_2_, 49 rhMG53 + H_2_O_2_, and 59 BSA + H_2_O_2_. (**C**) HEDA oxidative stress induced release of ROS from the mitochondria, as indicated with an increase in MitoSOX intensity after HEDA treatment. Treating HL-1 cells with rhMG53 significantly reduced mitochondria-generated ROS in the cytoplasm. (**D**) MitoSOX intensity was quantified as the integrated density for each cell. N=281 BSS BSA, 191 BSS rhMG53, 440 HEDA BSA, and 252 HEDA rhMG53.

In parallel studies, we treated HL-1 cells with H_2_O_2_, as another model of oxidative damage, and assessed mitochondria membrane integrity. Mitochondria membrane integrity is determined by measuring the transmembrane potential, which should be high in intact mitochondria, using the dye tetramethylrhodamine ethyl ester (TMRE). Consistent with earlier studies on mouse hearts [9], rhMG53 prevented the loss of TMRE signal after oxidative injury caused by H_2_O_2_ that occurred when BSA was present (**Figure 3C and D**). Maintenance of mitochondrial transmembrane potential indicates MG53 can prevent injury to the mitochondrial membrane.

### rhMG53 binds to cardiolipin to preserve mitochondria integrity under oxidative stress

We performed subcellular fractionation of HL-1 cells with or without HEDA treatment to separate the cytosolic proteins from proteins associated with the mitochondria. As shown in **Fig. 4A**, MG53 was enriched in the mitochondria fraction after HEDA damage compared to the mitochondria fraction under normal conditions. The presence of only GAPDH in the cytosolic lanes and COX IV in the mitochondrial lanes of the western blot indicate clear separation of the subcellular fractions.

**Figure 4:**
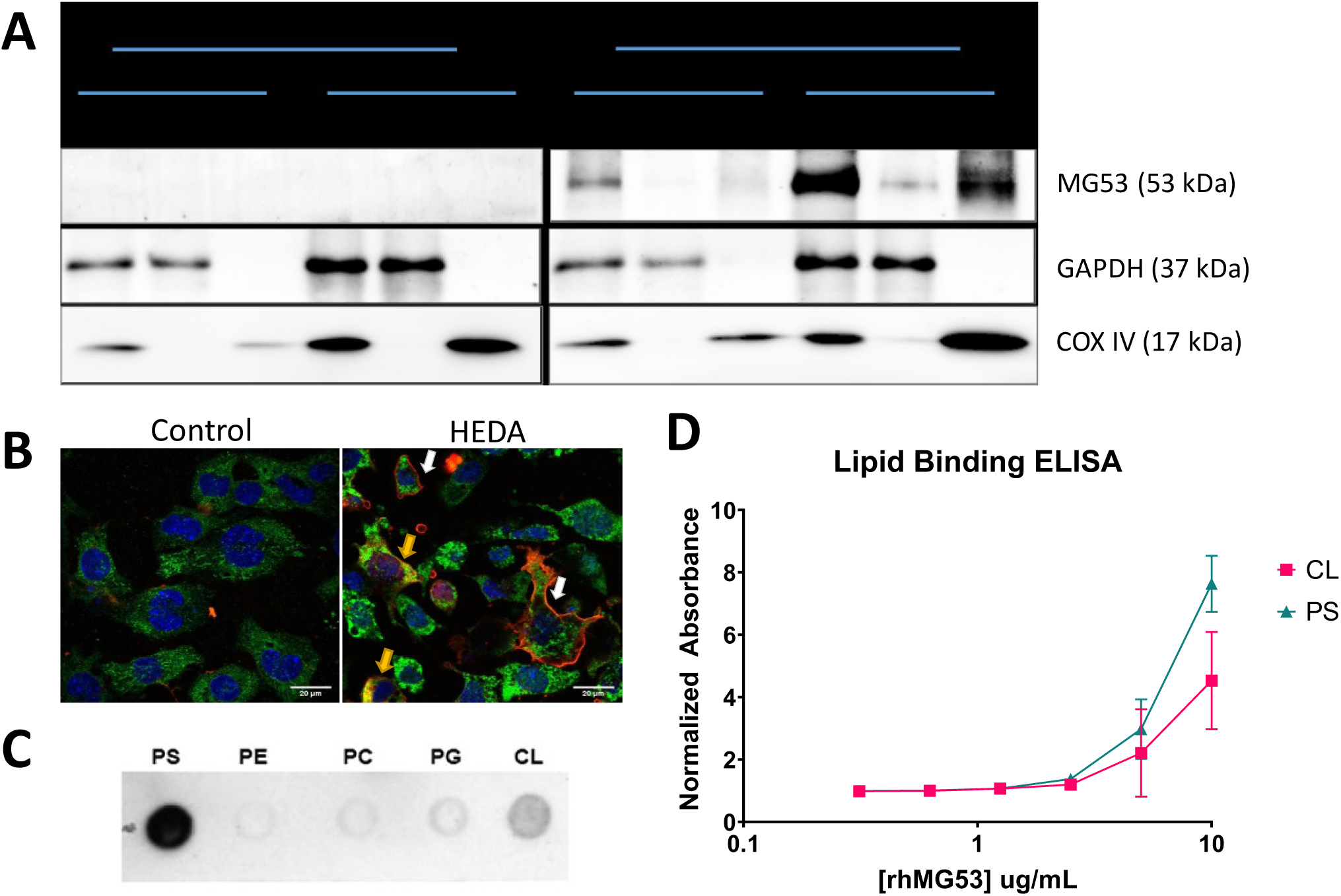
Oxidative stress leads to enrichment of rhMG53 at mitochondria in HL-1 cells; rhMG53 binds to mitochondria lipid cardiolipin. (**A**) HL-1 cells treated with either 10μg/mL BSA or rhMG53 underwent normoxia (BSS) or HEDA treatment. Western Blot analysis revealed an increased update of rhMG53 after HEDA damage and a strong co-localization of MG53 in the mitochondrial fraction. Purity of fractionation indicated with GAPDH and COX IV blotting. (**B**) rhMG53 binds to both phosphatidylserine (PS) and cardiolipin (CL). MG53 exhibits little or no binding to phosphatidylethanolamine (PE), phosphatidylcholine (PC), or phosphatidylglycerol (PG). (**C**) rhMG53 tagged with Alexa647 binds to both the plasma membrane (white arrow) and localize to mitochondria (yellow arrow) of HL-1 after HEDA treatment. (**D**) Quantitative lipid-based ELISA revealed dose-dependent binding of rhMG53 to PS and CL. N=4

Confocal imaging confirmed the results of the subcellular fractionation (**Figure 4B**). Cells that were not damaged did not have any rhMG53 entry into the cell. Conversely, damaged cells exhibited rhMG53 signal at both the plasma membrane (white arrows) and inside the cell surrounding the mitochondrial COX IV signal (yellow arrows).

We have previously shown rhMG53 directly interacts with the PS on the plasma membrane to facilitate protection against injury to tissues [4]. Thus, it is possible that MG53 may also be interacting with a phospholipid present on the mitochondria to facilitate mitochondria membrane repair injury during oxidative stress. We used a lipid dot-blot assay to test whether rhMG53 can bind to CL, a phospholipid that resides on the inner mitochondrial membrane that is flipped to the outer mitochondrial membrane after mitochondrial injury. The dot-blot assay shown in **Figure 4C** demonstrates that MG53 binds to PS as well as CL, but not other membrane phospholipids. Quantitative lipid-based ELISA revealed dose-dependent binding of rhMG53 to PS and CL (**Figure 4D**).

### rhMG53 prevents engulfment of mitochondria by lysosome after HEDA

Cells undergo mitophagy to preserve a pool of healthy mitochondria and degrade damaged mitochondria. To determine the extent to which MG53 is involved in the mitophagy process, HL-1 cells were infected with lentivirus carrying a mitochondrial-tagged keima fluorescent protein, mt-keima. mt-keima undergoes changes in the spectrum of fluorescence in the pH range from 3 to 8; fluorescing green at a neutral pH like the cytoplasm, and shifting to red under acidic pH, similar to that found in the lysosome [27].

**Figure 5** demonstrates HL-1 cells pre-treated with BSA that underwent oxidative damage showed a strong shift towards red fluorescence, an indicator of mitochondria presence in the lysosomes. Treatment with rhMG53 prevents that shift with mitochondria maintaining their bright green fluorescence. This suggests treating cells with rhMG53 reduces mitochondria encasement by the lysosome for degradation.

**Figure 5:**
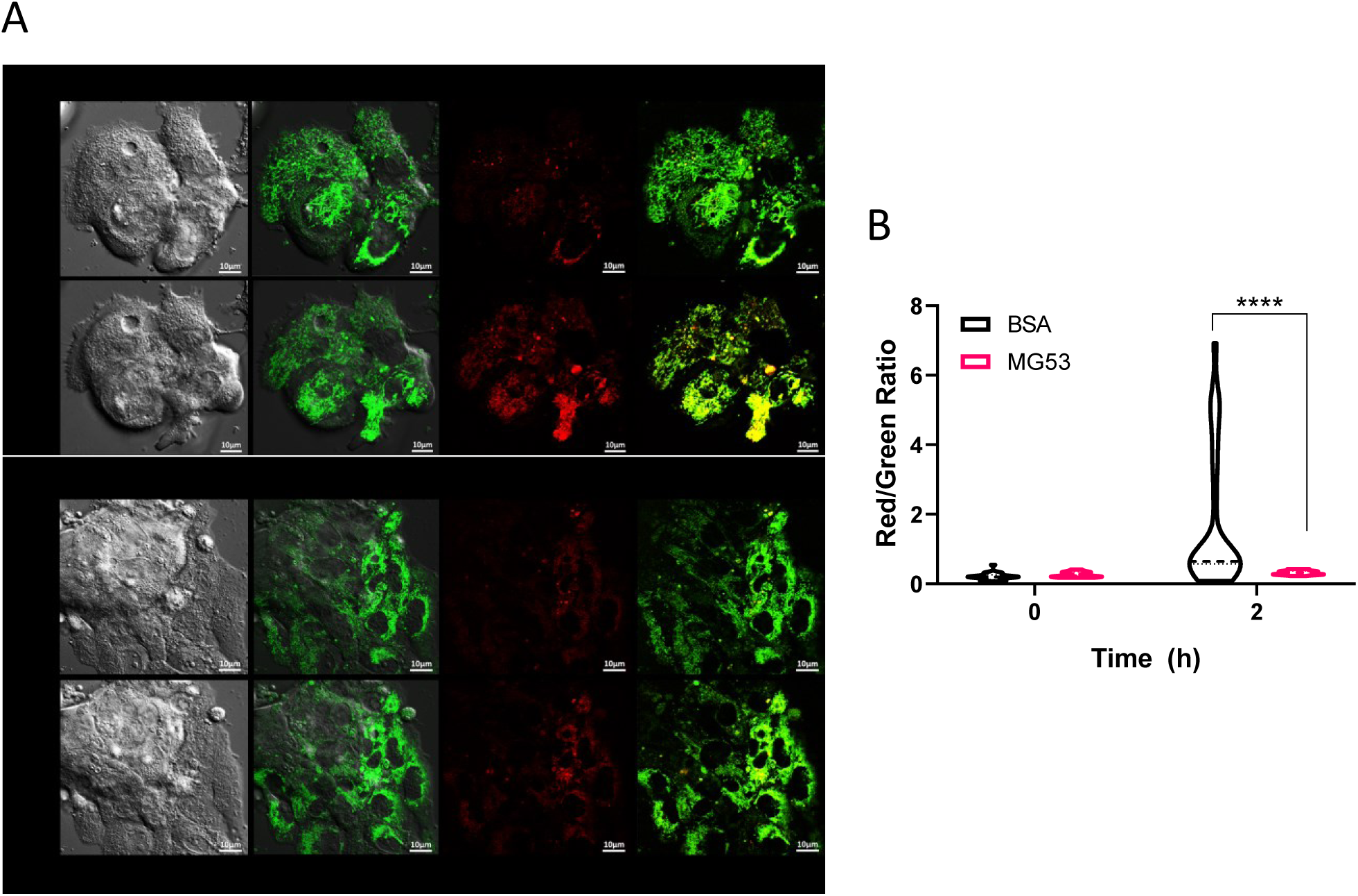
rhMG53 reduces lysosome-engulfment of mitochondria after H_2_O_2_ treatment in HL-1 cells. (**A**) HL-1 cells were infected with mt-Keima virus and underwent hypoxia/reoxygenation treatment after a 1hr pre-treatment of 10ug/mL BSA or rhMG53. Imaging revealed a strong shift towards red fluorescence indicating acidic pH in cells treated with BSA while cells treated with rhMG53 retained the green fluorescence of neutral pH. (**B**) Fluorescent intensity density for each cell was quantified and a ratio of red to green was calculated to determine degree of mitophagy present. Data was anlayzed via 2-way ANOVA (Prism Graphpad). N=28 BSA, and 16 rhMG53

## Discussion

Finding methods of tissue preservation after ischemic events like heart attack is essential to improving survival. Maintaining a healthy mitochondria population after ischemic insult is essential for cell, and consequently tissue, survival. We have previously described how MG53, either endogenous or exogenous, can protect cells and tissue from ischemic injury by fixing damage to the plasma membrane [8-11, 28, 29]. Here, we provide evidence that MG53’s ability to improve cardiomyocyte survival and function after ischemic injury is also linked to its ability to preserve a healthy pool of mitochondria.

We identified exogenous rhMG53 translocating to both plasma membrane and mitochondria after oxidative stress injuries like MI, H_2_O_2_, and HEDA. Co-localization of MG53 with the mitochondria was correlated with maintenance of mitochondrial membrane potential, and reduction in mitochondrial-associated ROS leak into the cytoplasm in both cultured HL-1 cells and primary adult mouse cardiomyocytes.

Previously, we have shown that MG53 can interact with PS, which is exposed from the inner leaflet of the plasma membrane upon damage, to facilitate tissue protection [30]. Here, we show MG53 can also bind to CL (**Figure 4**), a mitochondrial-specific lipid that is flipped from the inner mitochondrial membrane to the outer mitochondrial membrane after oxidative stress [31, 32]. This suggests MG53 may function similarly to preserve mitochondria integrity as it does to the plasma membrane.

Previous studies have shown that the human heart contains lower level of endogenous MG53 than rodent hearts, suggesting the need for exogenous application of the protein for cardioprotection [33]. Our present study demonstrates that rhMG53 has therapeutic value for mitochondrial maintenance as a means for cardioprotection. We provide biochemical and live cell imaging data to show that exogenous rhMG53 can enter the cells and target mitochondria to preserve their integrity under oxidative stress conditions. rhMG53 as a therapeutic agent is likely to be safe as the protein is naturally expressed in skeletal muscle and secreted into the serum as a myokine [5]. Although there is currently a debate regarding MG53’s role in insulin sensitivity via MG53’s ubiquitination of IRS-1 [13, 34-38], we have recently shown that sustained overexpression of MG53 does not alter metabolism while increasing regenerative capacity [5].

Since MG53 binds to CL on the outer mitochondrial membrane and results in a reduction of mito-ROS in the cytoplasm and maintenance of mitochondrial membrane integrity, we conclude that MG53 acts directly upon the mitochondria. Interestingly, the engulfment of mitochondria by lysosome, which is involved in mitophagy, was suppressed by rhMG53 treatment. This is not surprising as binding of rhMG53 to CL preserves integrity of the mitochondria, likely reducing the need for mitophagy.

Overall, this study provides a novel mechanism behind how rhMG53 treatment may be a clinically relevant strategy to reduce cardiomyocyte injury and maintain cardiac function in patients after ischemic injury. MG53’s function in preserving mitochondria function may contribute to preservation of the heart after ischemia/reperfusion injury. Understanding MG53’s interactions with mitochondria could be an attractive avenue for the development of MG53 as a targeted protein therapy for cardioprotection and potentially other regenerative medicine applications.

## References

1. Nowbar, A.N., et al., Mortality From Ischemic Heart Disease. Circ Cardiovasc Qual Outcomes, 2019. 12(6): p. e005375.

2. Liu, F., et al., Mitochondria in Ischemic Stroke: New Insight and Implications. Aging Dis, 2018. 9(5): p. 924–937.

3. Sucher, R., et al., Intracellular signaling pathways control mitochondrial events associated with the development of ischemia/reperfusion-associated damage. Transpl Int, 2009. 22(9): p. 922–30.

4. Cai, C., et al., MG53 nucleates assembly of cell membrane repair machinery. Nat Cell Biol, 2009. 11(1): p. 56–64.

5. Bian, Z., et al., Sustained elevation of MG53 in the bloodstream increases tissue regenerative capacity without compromising metabolic function. Nat Commun, 2019. 10(1): p. 4659.

6. Hwang, M., et al., Redox-dependent oligomerization through a leucine zipper motif is essential for MG53-mediated cell membrane repair. Am J Physiol Cell Physiol, 2011. 301(1): p. C106–14.

7. Weisleder, N., et al., Recombinant MG53 protein modulates therapeutic cell membrane repair in treatment of muscular dystrophy. Sci Transl Med, 2012. 4(139): p. 139ra85.

8. Liu, J., et al., Cardioprotection of recombinant human MG53 protein in a porcine model of ischemia and reperfusion injury. J Mol Cell Cardiol, 2015. 80: p. 10–19.

9. Wang, X., et al., Cardioprotection of ischemia/reperfusion injury by cholesterol-dependent MG53-mediated membrane repair. Circ Res, 2010. 107(1): p. 76–83.

10. Duann, P., et al., MG53-mediated cell membrane repair protects against acute kidney injury. Sci Transl Med, 2015. 7(279): p. 279ra36.

11. Jia, Y., et al., Treatment of acute lung injury by targeting MG53-mediated cell membrane repair. Nat Commun, 2014. 5: p. 4387.

12. Chung, Y.W., et al., Targeted disruption of PDE3B, but not PDE3A, protects murine heart from ischemia/reperfusion injury. Proc Natl Acad Sci U S A, 2015. 112(17): p. E2253–62.

13. Ma, H., et al., Effect of metabolic syndrome on mitsugumin 53 expression and function. PLoS One, 2015. 10(5): p. e0124128.

14. Lijie, G., et al., Mitsugumin 53 promotes mitochondrial autophagy through regulating Ambra1 expression in C2C12 myoblast cells. Cell Biol Int, 2019. 43(3): p. 290–298.

15. Fang, H., et al., Imaging superoxide flash and metabolism-coupled mitochondrial permeability transition in living animals. Cell Res, 2011. 21(9): p. 1295–304.

16. Ding, Y., et al., Mitoflash altered by metabolic stress in insulin-resistant skeletal muscle. J Mol Med (Berl), 2015. 93(10): p. 1119–30.

17. Wang, Z., et al., Irisin Protects Heart Against Ischemia-Reperfusion Injury Through a SOD2-Dependent Mitochondria Mechanism. J Cardiovasc Pharmacol, 2018. 72(6): p. 259–269.

18. Xiao, Y., et al., ROS-related mitochondrial dysfunction in skeletal muscle of an ALS mouse model during the disease progression. Pharmacol Res, 2018. 138: p. 25–36.

19. Wang, H., et al., ALS-associated mutation SOD1(G93A) leads to abnormal mitochondrial dynamics in osteocytes. Bone, 2018. 106: p. 126–138.

20. Zhou, J., et al., Dysregulated mitochondrial Ca(2+) and ROS signaling in skeletal muscle of ALS mouse model. Arch Biochem Biophys, 2019. 663: p. 249–258.

21. Schindelin, J., et al., Fiji: an open-source platform for biological-image analysis. Nat Methods, 2012. 9(7): p. 676–82.

22. Claycomb, W.C., et al., HL-1 cells: a cardiac muscle cell line that contracts and retains phenotypic characteristics of the adult cardiomyocyte. Proc Natl Acad Sci U S A, 1998. 95(6): p. 2979–84.

23. White, S.M., P.E. Constantin, and W.C. Claycomb, Cardiac physiology at the cellular level: use of cultured HL-1 cardiomyocytes for studies of cardiac muscle cell structure and function. Am J Physiol Heart Circ Physiol, 2004. 286(3): p. H823–9.

24. Astrom-Olsson, K., et al., Impact of hypoxia, simulated ischemia and reperfusion in HL-1 cells on the expression of FKBP12/FKBP12.6 and intracellular calcium dynamics. Biochem Biophys Res Commun, 2012. 422(4): p. 732–8.

25. Lazarou, M., et al., The ubiquitin kinase PINK1 recruits autophagy receptors to induce mitophagy. Nature, 2015. 524(7565): p. 309–314.

26. Wang, W., et al., Superoxide flashes in single mitochondria. Cell, 2008. 134(2): p. 279–90.

27. Sun, N., et al., A fluorescence-based imaging method to measure in vitro and in vivo mitophagy using mt-Keima. Nature Protocols, 2017. 12(8): p. 1576–1587.

28. Cao, C.M., et al., MG53 constitutes a primary determinant of cardiac ischemic preconditioning. Circulation, 2010. 121(23): p. 2565–74.

29. Yao, Y., et al., MG53 permeates through blood-brain barrier to protect ischemic brain injury. Oncotarget, 2016. 7(16): p. 22474–85.

30. Nagata, S., et al., Exposure of phosphatidylserine on the cell surface. Cell Death Differ, 2016. 23(6): p. 952–61.

31. Paradies, G., et al., Role of Cardiolipin in Mitochondrial Function and Dynamics in Health and Disease: Molecular and Pharmacological Aspects. Cells, 2019. 8(7).

32. Paradies, G., et al., Role of cardiolipin peroxidation and Ca2+ in mitochondrial dysfunction and disease. Cell Calcium, 2009. 45(6): p. 643–50.

33. Lemckert, F.A., et al., Lack of MG53 in human heart precludes utility as a biomarker of myocardial injury or endogenous cardioprotective factor. Cardiovasc Res, 2016. 110(2): p. 178–87.

34. Song, R., et al., Central role of E3 ubiquitin ligase MG53 in insulin resistance and metabolic disorders. Nature, 2013. 494(7437): p. 375–9.

35. Hu, X. and R.P. Xiao, MG53 and disordered metabolism in striated muscle. Biochim Biophys Acta Mol Basis Dis, 2018. 1864 (5 Pt B): p. 1984–1990.

36. Yi, J.S., et al., MG53-induced IRS-1 ubiquitination negatively regulates skeletal myogenesis and insulin signalling. Nat Commun, 2013. 4: p. 2354.

37. Liu, F., et al., Upregulation of MG53 induces diabetic cardiomyopathy through transcriptional activation of peroxisome proliferation-activated receptor alpha. Circulation, 2015. 131(9): p. 795–804.

38. Wu, H.K., et al., Glucose-Sensitive Myokine/Cardiokine MG53 Regulates Systemic Insulin Response and Metabolic Homeostasis. Circulation, 2019. 139(7): p. 901–914.

